# Model recapitulates regenerative limb blastema formation through local softening of the wounded epithelium

**DOI:** 10.64898/2026.03.11.711112

**Authors:** Samantha Finkbeiner, Ansa Brew-Smith, Xueqing Wang, Dzi Tsiu Fu, James Monaghan, Calina Copos

## Abstract

Studies of appendage regeneration in vertebrates have shown that the fundamental building block of any regenerative tissue is a blastema. The blastema is a cone-shaped accumulation that forms at the site of amputation post wound-healing and is the result of a highly coordinated process involving a cluster of cells capable of growth, migration, and differentiation. Although several key signaling pathways involved in regeneration have been identified, which cellular processes they control and how these processes are coordinated across space and time are not yet fully understood. This study introduces a computational tool to examine how the outgrowth results from the interaction of two tissue layers: the bulk (mesenchyme) and the overlying epithelium. We developed a novel hybrid agent-based modeling framework and an accompanying parameter inference pipeline to uncover the cellular properties in the epithelium and the mesenchyme driving the formation of a normal regenerative blastema with a morphology similar to that observed in experiments. Using our model, we report two conditions for blastema formation: retained local softening of the epithelial layer at the site of injury, which was confirmed experimentally with atomic force microscopy (AFM) measurements, and the involvement of the Wnt signaling pathway in the directed migration of mesenchyme cells towards the distal tip. Taken together, this combined experimental-theoretical approach provides a framework for understanding how the Wnt signaling pathway influences the formation of the early blastema at multiple levels of organization and how key cellular behaviors contribute to its formation.

**Author Summary:** A small number of tetrapods have retained the ancestral ability to regenerate tissues and even limbs. Indifferent of species or tissue, the decisive initial stage of limb regeneration is the formation of a specialized structure called the blastema, a heterogeneous mass of mesenchymal cells, in a relatively short timescale of 2-14 days post injury or amputation. To study the mechanical and cellular conditions for limb blastema formation in the axolotl model organism, we developed a novel hybrid agent-based modeling framework and accompanying kinetic parameter inference pipeline. By recapitulating blastema morphometrics of healthy and stalled regenerative states, our model finds two conditions for blastema formation: retained local softening of the epithelial layer at the injury site post wound-healing, which we confirmed with atomic force microscopy measurements, and that the Wnt signaling pathway plays a role in the migration of mesenchyme cells to the distal tip in order to produce the blastema.

## INTRODUCTION

Axolotls (*Ambystoma mexicanum*) are valuable vertebrate models for studying regeneration, capable of regenerating complex structures, including limbs, the heart, spinal cord, and other organs [1, 2] (Fig. 1A). During the first weeks after limb amputation, a conserved molecular toolkit (FGF, Wnt, BMP, Neuregulin, Hedgehog, retinoic acid) coordinates cellular dynamics to establish a blastema — a critical tissue that distinguishes an appendage (and therefore, an organism) that can and cannot regenerate [3]. The blastema is a cone-shaped outgrowth covered by a rigid protective skin layer called the epithelium. Its interior consists of the mesenchyme, a heterogeneous collection of cells derived from tissues just proximal to the amputation site [4–6]. This regenerative outgrowth is established as early as 5 days post amputation (dpa) through a highly coordinated process; it starts with wound closure in the wounded epithelium within 12-24 hours, driven by the migration of nearby epithelial cells, innervation, cell de-differentiation and migration, proliferation, formation of a specialized signaling center called an apical epithelial cap (AEC), and finally the onset of differentiation (Fig. 1B). Removal of the AEC blocks regeneration, and transplantation of an ectopic AEC on an early blastema leads to an ectopic blastema, demonstrating it is necessary for the early phase of limb regeneration [7, 8]. Over the next 5-7 days, cells primarily of connective tissue fibroblast origin aggregate under the AEC, generating the blastema with a similar structure and gene expression to that of an embryonic limb bud. Despite considerable efforts to understand the cellular makeup and cell-cell signaling required to generate a blastema, there is a gap in knowledge about how highly organized cells at the limb stump reorganize into a seemingly homogeneous blastema [9].

**Figure 1.**
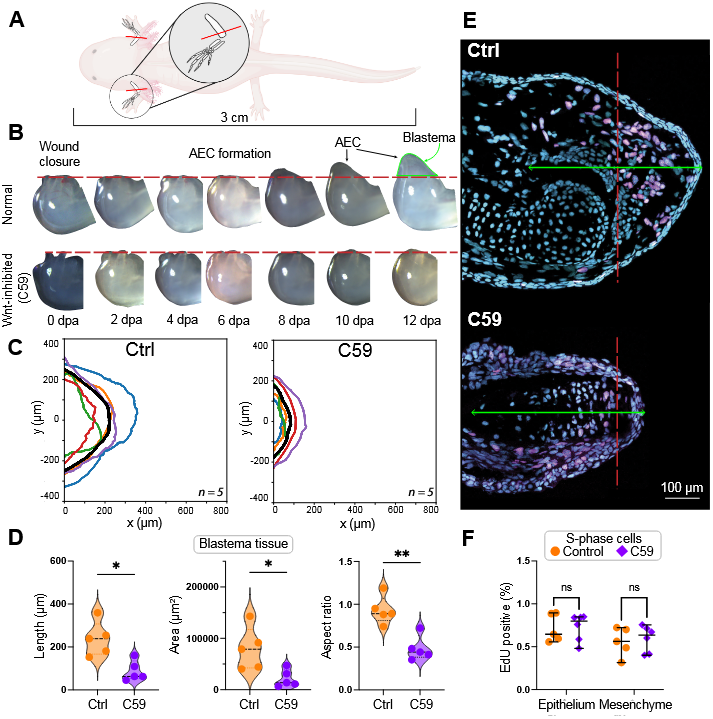
Quantitative analysis of 2D cross-sections of blastema tissue and nuclear morphometrics of control and Wnt-inhibited regenerating limbs at 7 dpa. (A) Schematic of axolotl and bilateral limb amputation assay. Red line, amputation plane. (B) Scheme of the phases and transitions of limb regeneration including apical epithelial cap (AEC) formation across 12 days post amputation (dpa). (C) Average outgrowth shape at 7 dpa in CTRL (left) and C59 (right). Thick black outlines represent averages, while colored outlines mark individual tissue samples. (D) Quantification of length, area, and aspect ratio of the outgrowth blastema at 7 dpa in CTRL and C59. (E) Representative histological blastema tissue samples at 7 dpa with EdU+ (S-phase) cells marked in magenta. Red dashed line, amputation plane; green line, 500 *μ*m from distal tip; scale bar: 100 *μ*m; tissue thickness: 16 *μ*m. (F) Quantification of S-phase cells in the blastema in the epithelium layer and mesenchyme in each condition. *N* = 5 animals/condition except in (F) where *N* = 6 animals were used for C59. Mann-Whitney test, * *p<* 0.05.

The Wnt signaling pathway is a promising therapeutic target for promoting blastema formation because it is necessary for appendage regeneration (and development) in many species, including Xenopus tails, zebrafish fins, salamander limbs, mammalian digit tips, and deer antler regeneration [10–17]. Several hypotheses regarding blastema outgrowth have been proposed so far [18]. Substantial data in other vertebrates suggest “growth-based morphogenesis,” hypothesizing that the outgrowth is driven by a higher proliferative rate in the limb-forming region. However, this mechanism would require proliferation rates that far exceed realistic values [19–21]. The study of limb development in mice and chicks has provided several models for limb outgrowth, including directional cell movement regulated by non-canonical Wnt signaling [19, 22, 23]. Mechanical properties such as phase separation of more fluidized limb cells from the flanking cells [24–29], or convergent extension by directed cell intercalation [30, 31] have been shown to be critical in the regulation of limb bud elongation in development.

Computational modeling of developing tissues and organs has served an important role in testing and posing hypotheses for both growth [31–35] and patterning [36,37]. Among them, agent-based models (ABM) are particularly useful in understanding how individual dynamics, heterogeneity, and local interactions influence emergent behavior. However, few models exist in the context of regeneration, and even fewer focus on blastema formation, the first and pivotal stage of regeneration. In an ABM, cells are represented as discrete particles, each with their own set of biological and physical attributes. The collective behavior of each individual allows one to explore hypotheses relating local changes to behaviors (proliferation, migration, etc.) at the population or community level.

In this study, we develop a hybrid two-dimensional (2D) model combining deterministic and stochastic ABM approaches to explore the contributions of individual cell morphometrics and mechanics to the formation of an accumulation blastema in control and Wnt-inhibited limbs. The model explicitly incorporates a variety of processes enabling the stochastic simulation of proliferating and migrating cells starting from a flat multilayer limb stump. Major model components include a nonuniform, deformable tissue layer that represents the surrounding epithelium; distinct cell cycle stages; implicit, continuous biochemical environments that inform biological and physical processes; and integrated biological and physical processes that govern cellular behaviors, including proliferation, apoptosis, cell-cell adhesion, migration, and deformation.

By recapitulating the morphology of control and Wnt-inhibited regenerative blastemas in simulations, our modeling framework makes two important biological predictions: (1) the presence of local softening of the epithelial layer post-wound healing, and (2) the distal-oriented persistent migration of mesenchymal cells towards the AEC underlies the outgrowth, which is controlled by the Wnt pathway. Atomic force microscopy (AFM) measurements confirm the mechanical softening of the wounded epithelium compared to the uninjured tissue at 7 dpa in both control and Wnt-inhibited regenerating limbs.

In addition to forward modeling, we developed a parameter inference pipeline to estimate key kinetic rates, including total cell cycle duration, persistent migration rate, and the fraction of cells undergoing directed migration. By minimizing the error between simulated and experimental blastema shapes, the optimization procedure identified a parameter manifold, indicating a trade-off and identifiability constraint between cell migration and proliferation. When the average duration of the cell cycle is constrained to experimentally reported values (40-53 hours [18]), the model predicts that at least 30% of cells undergo persistent migration toward the AEC with an average speed of 50-74 μm/day. For Wnt-inhibited limbs, the inference procedure predicts either a markedly prolonged cell cycle (exceeding 240 hours) or the loss (or substantial reduction) of AEC-directed mesenchymal cell migration. Because our experimental data do not indicate reduced proliferation under full Wnt inhibition, we conclude that Wnt signaling regulates distal-directed persistent migration of mesenchymal cells towards the AEC. Taken together, these combined experimental and theoretical approaches establish a framework for identifying the cellular dynamics governing blastema formation in normal limb regeneration and provide insight into the dysregulated processes that arise when Wnt signaling is impaired.

## MODEL OVERVIEW

We model the composite blastema tissue within the well-established agent-based framework. This coarsegrained framework represents cells as interacting rigid particles in a longitudinal two-dimensional (2D) slice through the middle of the blastema tissue, 600 *μ*m wide and 400 *μ*m long before the amputation plane (Fig. 2A-B). Because the experiments visualize the outgrowth tissue and cellular dynamics in a cross-section of the limb bud, we similarly restrict our simulations to a 2D plane. The *x* and *y* axes in the computational domain represent the proximodistal (PD) and anteroposterior (AP) anatomical axes, respectively (Fig. 2B). We ignore the effects of bone and cartilage and instead focus on the cells in the mesenchyme enclosed by a deformable epithelium.

**Figure 2.**
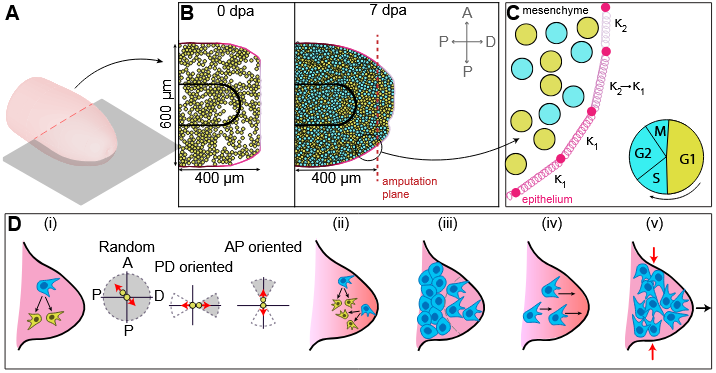
Diagram of the computational model and possible hypotheses for early regenerative blastema formation. (A) 3D schematic of regenerating limb and the 2D cross-section representing the computational domain. (B) Realization of the modeling framework at 0 and 7 dpa. Inset, anatomical axes: horizontal (proximodistal, PD) and vertical (anteroposterior, AP). (C) Zoom-in of the model showing that the 1D deformable structure of the epithelium is described as elastic, prestress springs connected in series with spatially variable stiffness. Cells in the mesenchyme are colored by their cell cycle state. Inset, cell cycle schematic. (D) Hypotheses for outgrowth: (i) oriented division and possible orientations of daughter cells following a cell division event, (ii) proliferation gradient, (iii) phase separation, (iv) directed migration, and (v) convergent intercalation.

Each mesenchymal cell is represented as a 2D object approximated by its center-of-mass, **x**_M_ representing an (*x, y*) coordinate pair, and its fixed size diameter *d*_0_. Details of the cell’s morphology, including membrane protrusions and cytoplasm, are ignored in this coarse-grained modeling approach. Each cell has a collection of attributes that describe its key biological and physical characteristics — these include position, velocity, cell type, and cell cycle information. The epithelial layer is coarse-grained into a homogeneous structure of nodes **x**_E_ interconnected by elastic, contractile springs. The body proximal regions of the epithelium are physically anchored in space with zero velocity. Mechanical heterogeneity of the structure is considered at a later stage. Epithelial and mesenchymal cells are mechanically coupled through a distance-dependent steric repulsion. This interaction is symmetric: when a mesenchymal cell exerts force on the epithelium (for instance, by randomly moving closer), the epithelium exerts an equal and opposite force back on the cell. The strength of this repulsive force has a cutoff distance.

### Cell cycle process

Our model employs a simplified description of the cell cycle with two phases: mitotic (S/G2/M) and at rest (G1), adapted from [38] and applied only to the mesenchyme tissue. Each mesenchymal cell begins in the G1 phase. The durations of the G1 and S/G2/M phases are modeled as geometrically distributed random variables with mean lengths T_G1_ and T_S*/*G2*/*M_, respectively. At each discrete time step, a cell transitions to the next phase with the following probabilities:

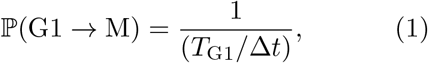

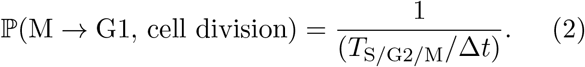

Cells have a constant probability of apoptosis at each time step:

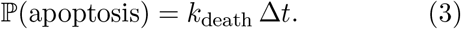

These probabilities are sampled independently at each time step. The S/G2/M *→* G1 transition results in cell division, with a daughter cell positioned in the simulation next to the mother at a random angle θ chosen from a uniform distribution (Fig. 2D), unless a different hypothesis is investigated. In the limit of large numbers, we expect the averaged behavior of the stochastic dynamics to converge to the following compartmental coupled differential equations:

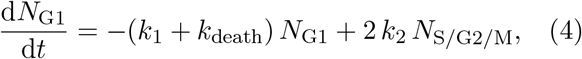

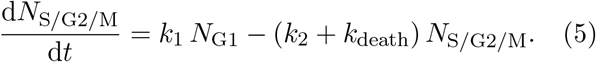

The number of cells in the G1 and S/G2/M phases are N_G1_ and N_S*/*G2*/*M_, respectively. The transition rates are defined as k_1_ = 1/T_G1_, k_2_ = 1/T_S*/*G2*/*M_, and k_death_, and the apoptosis rate is k_death_. We numerically solved this system of differential equations and found good agreement with the averaged discrete behavior of individual cells.

### Physical processes

The physical processes describe the movement of mesenchymal cells and the deformation of the epithelium. Taking the overdamped (or viscous dominated) approximation to a physical system [39, 40], each structure moves with its own velocity according to local force density balance equations:

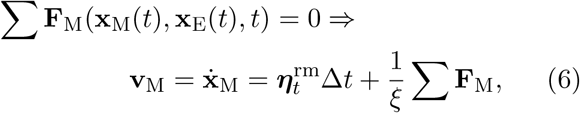

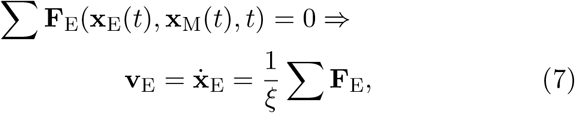

where ξ is the viscous drag coefficient, effectively equivalent to elastic cell-matrix interactions under conditions of high dissociation rates of these adhesion bonds [41]. Forces acting on each structure are defined below.

### Cells in the mesenchyme

The force densities on a mesenchymal cell *i* are

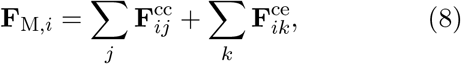

where 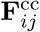 is the cell-cell interaction between neighboring mesenchymal cells *i* and *j*, and 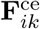 is the pairwise force density exerted on mesenchymal cells and epithelium nodes to resolve overlap. 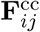 follows a simple force law:

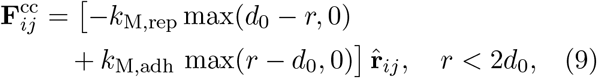

namely, repulsive on short length scales and attractive on longer length scales as outlined in [42, 43]. We define **r**_*ij*_ = **x**_*j*_ *-* **x**_*i*_ as the unit directional vector pointing from cell *i* to cell *j*. This force is zero outside the cutoff distance of 2d_0_ to reduce the computational cost using a parallel cell-list algorithm (SI, Note A). The force 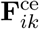 takes the same form as in Eq. (9), but with a stronger repulsion constant (k_ME,adh_ > k_M,adh_). An equal and opposite reactive force is applied to the epithelium node k as reflected in Eq. (10). Lastly, mesenchymal cells undergo Brownian motion with displacement incre-ments: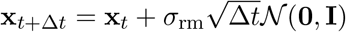 .

### Epithelium

The force density at a material node *k* along the epithelial structure **x**_E_ is

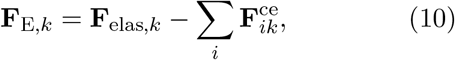

where **F**_elas_ is the elastic force density computed as

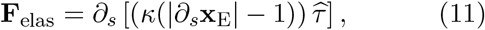

with an elastic response of stiffness *κ* (units of force/length, Fig. 2B). Here, s is the local parametric coordinate on the epithelium boundary, and 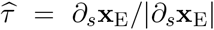 is a unit vector tangent to the boundary. For simplicity, proliferation in the epithelial layer is ignored, which is supported in the literature [18, 44].

## RESULTS

### Wnt signaling is required for limb regeneration

To identify the role of Wnt signaling in the formation of an accumulation blastema during limb regeneration in axolotls, we performed Wnt-inhibition experiments using a PORCN inhibitor to block the secretion of all Wnt ligands. This method of Wnt inhibition has been used previously to show that the inhibition of Wnt signaling stops both development and regeneration in juvenile axolotls [10], but the changes in tissue and cell morphometrics were not detailed in these studies. Similarly, we found a significant decrease in both blastema area and length in the Wnt inhibited treatment group compared to controls (Fig. 1C-D). The Wnt inhibited treatment group showed wound closure and few cells between the skeletal elements and the epithelium, rather than an accumulation of cells that is present in normal regeneration. These differences in outgrowth at 7 dpa resulted in different tissue shapes between the treatment groups. The Wnt inhibited group had a flatter distal end of the limb compared to the cone-shaped normal blastema (Fig. 1D). A normal blastema at 7 dpa has an average area of 0.09 mm^2^, an average length of 240 μm, and an aspect ratio close to 1, indicating that the outgrowth length is close to the width of the amputated limb. By contrast, the Wnt-inhibited treatment group has an average blastema area of 0.02 mm^2^, roughly one quarter the size of a normal blastema, a smaller average length of 90 μm, and an aspect ratio just below 0.5. Given these tissue-level growth differences, we next examined whether corresponding differences in cellular morphology, structural organization, or dynamics were present

Next, we characterized the morphologies of individual nuclei during normal regeneration, starting 500 μm proximal to the distal tip of the limb at 7 dpa, which has been experimentally observed in previous studies [45–47] (Fig. 1E). Staining of the nucleus and plasma membrane provides morphological information in the stained sections. No significant changes in area, aspect ratio, and circularity were noted overall when comparing the Wnt inhibited treatment to controls (Fig. S1A-C).

The number and spatial location of proliferating cells during limb regeneration were analyzed using EdU staining, which marks the S phase of the cell cycle. We did not note a difference in the total percentage of cells that were EdU+ between the two conditions, either in the whole tissue, between the two tissue layers (Fig. 1F), or when further quantified spatially by 100 μm sections (Fig. S1D). These results lead us to infer that proliferation is not the main driving factor behind the outgrowth of an accumulation blastema, and that similarly, the cellular morphological quantification provides limited information about underlying processes.

### A computational model for limb regrowth from a flat tissue

To dissect which cellular factors contribute to the initial cone-shaped bud outgrowth, we attempt to recapitulate the phenomenon using a hybrid agent-based modeling framework (Fig. 2, Movie 1). The model is initialized by randomly distributing 500 mesenchyme cells within a 2D cross-section of the limb stump 600 *μ*m wide and 400 *μ*m long before a flat amputation plane (Fig. 2A-B). The mesenchyme cells are confined to the region by a deformable epithelial tissue layer (Fig. 2C) with homogeneous stiffness *κ*, except at the anterior and posterior regions where stiffness is twice as large to enforce attachment to the animal body. The system evolves according to a viscous force density balance from Eqs. (6)-(7), along with stochastic biological events of cell proliferation and apoptosis. The ensemble cell cycle dynamics are approximated by an equivalent system of differential equations (Eqs. (4)-(5), Fig. S2). Equations of motion are solved numerically using the Forward Euler-Maruyama integration scheme (SI, Note A), and parameter values are provided in Table S1. In SI Note B, a justification of parameter values is provided.

Fig. 2B plots the resulting outgrowth shape — mesenchyme cells randomly migrate with an effective diffusion coefficient of 0.5 *μ*m^2^/day and also undergo proliferation with an average cell cycle of 56 hours. Along with the outgrowth shape (star, Fig. 3), we also plot the time-evolution of the final cellular density and effective volume fraction (star, Fig. 3E, Fig. S3). We find that the simulated outgrowth length, area, and aspect ratio are well below what we observed in control cases at 7 dpa (star, Fig. 3A-F). This means that our model, without additional assumptions, cannot capture the observed regenerative bud formation in control or even in Wnt-inhibited regenerating limbs.

**Figure 3.**
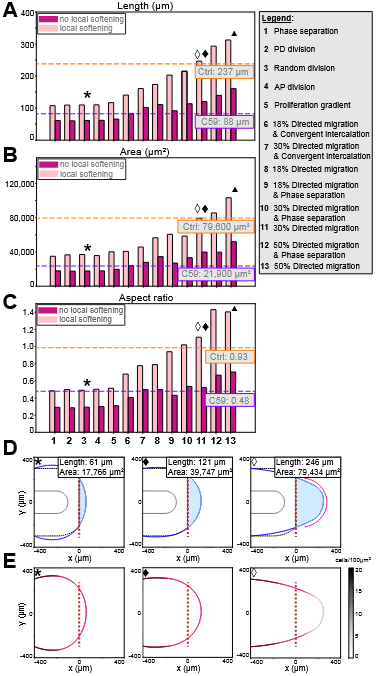
Comparison of simulated outgrowth morphology with and without local epithelium softening for the stated hypotheses of early regeneration. Morphological measurements include: (A) length (*μ*m) (B) area (*μ*m^2^), and (C) aspect ratio. Outgrowth length is measured as the maximum distance along the PD axis. The horizontal axis is labeled by the tested hypotheses. For sample simulation realization (D) the outgrowth shape and (E) cellular density are shown at 7 dpa. Star, default setup of random oriented proliferation with no local softening or additional motility assumptions. Diamond and triangle, 30% and 50% of mesenchymal cells migrate directionally, respectively.

We know that a variety of signaling morphogens, including Wnt, are implicated in tissue regeneration (and development). In this framework, we test the model’s response to a variety of hypotheses that could lead to a more sizable outgrowth, including: (i) preferential orientation of the division plane along the PD or AP axes, (ii) enhanced proliferation along a spatial gradient, (iii) local phase separation, (iv) directed migration in the distal direction, and (v) convergent extension-like intercalation (Fig. 2D). Briefly, we describe the biological assumptions implemented for each hypothesis; additional details are provided in SI, Note C.

i. Preferentially oriented division: During cell division, the position of the daughter cell relative to the mother cell is determined by the orientation of the division angle. These angles follow normal distributions: (i) for PD divisions, the mean is 0 radians (SI, Note C, Eq. S1), while for AP divisions, the mean is π/2 radians (SI, Note C, Eq. S2). Both distributions have a standard deviation of π/6 radians.
ii. Proliferation gradient: A spatial proliferation gradient is created by modulating the G1 phase duration in mesenchyme cells. As a regulatory boundary moves proximally from the distal tip 50 μm per day, cells located distal to this boundary experience shortened G1 phases from 28 to 18 hours, while cells positioned proximal to the boundary undergo lengthened G1 phases from 28 to 36 hours. This results in a time-to-division gradient with faster proliferation in the distal tip compared to the body proximal end.
iii. Phase separation: A phase-separated tissue architecture is established with a more solid-like proximal confluent region and a more fluid-like distal zone in the mesenchyme tissue. Specifically, mesenchyme cells outside a region 200 μm from the distal tip are in a confluent, solidlike state enforced through zero random motion 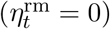 .
iv. Directed migration: Similar to (ii), a motility gradient is created by introducing biased motility in mesenchymal cells. As the regulatory boundary moves from the distal tip proximally by 50 μm each day, a fixed percentage of cells located distal to this boundary experience a biased motion term 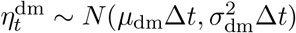 .
v. Convergent intercalation: Similarly, a biased motion term is added to a fixed percentage of mesenchymal cells, but with two notable differences: (1) the bias is towards the midline in the AP axis, and (2) a fixed percentage of cells on either side of the midline are subjected to this motion. Intercalating cells move according to the same mean and variance as those of directed migration in (iv).

In addition to these spatial dynamics, the percentage of cells responding to these morphogen-altered dynamics is also interrogated. Under the assumptions of each hypothesis, we quantify the length, area, and aspect ratio of the outgrowth past the amputation plane (Fig. 3, Fig. S4). Regardless of which hypothesis is probed, the simulated outgrowth is underestimating the actual outgrowth in control cases. For example, even with 50% of the cells migrating directionally at a speed of 100 *μ*m/day, by 7 dpa, the simulated blastema reaches a length that is roughly 66% of the physical regenerative bud (Fig. 3A-C, triangle).

### Model posits local softening of the wounded epithelium in the early regenerative blastema

Next, we ask whether varying the mechanical response of the epithelial layer could improve the model’s predicted outgrowth. Specifically, we focus on stiffness, the passive mechanical property of an object to deform under load. To enforce this, we implemented a stiffness patterning along the epithelium at all time points — at the distal tip, the stiffness coefficient is lowered roughly 100-fold in a region 400 *μ*m centered at the midline (Fig. 2B). Under the assumption of local epithelial softening, two hypotheses reproduce outgrowths of a similar or larger size and aspect ratio as observed: *directed motility* and *phase separation with directed motility*. Based on our findings, at least 30% of cells exhibit biased migration toward the distal tip at any given time point, and if a phase-separated tissue architecture is present, at least 40% of cells within that distal zone demonstrate distal-biased migration (Movie S2). Model simulations without softening (purple) report approximately half the outgrowth length as those with local softening (pink), as shown in Fig. 3.

To investigate whether a spatial gradient of mechanical softening is indeed required, we revisited these hypotheses in the presence of an overall softer epithelial layer. Specifically, the overall epithelial stiffness was similarly reduced. Consistently, we found that the model produced a shorter, rounder outgrowth rather than the expected cone-shaped blastema (Fig. S5A). Even spatial broadening, mis-localization, or mechanical heterogeneity of the softer epithelium region did not reproduce the expected outgrowth (Fig. S5B-C). Thus, we conclude that local softening of the epithelium is necessary to generate a blastema similar in size to those of control limbs. Next, we determine whether this local softening of the epithelium is biologically relevant.

### AFM measurements confirm local epithelial softening in regenerating limbs

To test the idea that the wounded epithelium remains mechanically soft long after wound closure, we performed atomic force microscopy to determine the apparent elastic moduli (hereafter referred to as stiffness) of regenerating tissues (Fig. 4). We directly measured the epithelial layer overlying the blastema (Fig. 4A). AFM analysis revealed a steep decrease in tissue stiffness in the wounded site compared to the intact site at 7 dpa, after wound closure had been completed (Fig. 4B). The observed tissue softening in the epithelial layer correlates with the establishment of prospective bud tissue, suggesting that softening enables the distal outgrowth of the underlying

**Figure 4.**
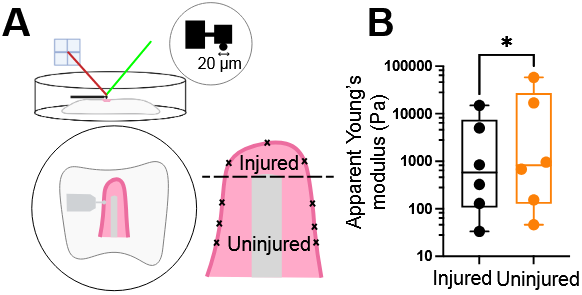
Stiffness measurements show local softening of the injured epithelium. (**A)** Limbs were collected at 7 dpa and sectioned with a vibratome for AFM measurements. (B) Apparent Young’s modulus values measured in the epithelium by AFM indentation using a cell-sized indenter (20 *μ*m diameter). The positions marked “x” indicate the approximate locations where AFM measurements were performed on the injured and uninjured epithelium. Dashed line, amputation plane. *N* = 6 animals/condition. Wilcoxon paired test, * *p<* 0.05.

These results confirmed that the injured epithelial layer is softer compared to the surrounding tissue long after wound closure, and this softening allows the underlying mesenchyme to produce an outgrowth. A similar differential tissue response underpins organ shaping, such as the buckling of the chick gut [33]. Through this revised perspective, we re-examined our findings in Fig. 3 and found that hypotheses combining directed cell migration with a mechanically soft wounded epithelium successfully recapitulated the observed blastema size. These results inform us that the softening of the epithelium enables the formation of a blastema with a morphology similar to that which was experimentally observed through the directional migration of mesenchyme cells towards the distal tip. The confirmation of the theory-inspired hypothesis of epithelial softening validated our modeling approach, but it remains to be determined whether this link is causal and how it is controlled by Wnt signaling.

### The composite blastema tissue response undergoes overall mechanical stiffening

The apparent local softening of the epithelial layer was at odds with our expectation of seeing an overall stiffening of the tissue during regeneration due to the increase in cellular density caused by proliferation. This led us to investigate the overall mechanical response of the whole-limb by simulating AFM-like poking experiments (Fig. 5A). For this setup, we use the directed migration hypothesis, with 30% of mesenchymal cells migrating directionally and local softening of the epithelium, as shown in Fig. 3 (hollow diamond). To do so, at each daily time point, we halt the dynamics of cell proliferation and apoptosis, as well as other dynamics from outgrowth hypotheses such as directed migration, intercalation, and phase separation. After allowing the friction-dominated system to reach mechanical equilibrium, a localized external force is applied to the distal tip for a constant time interval, emulating AFM measurements. The external force per unit length is constant across time points and is applied along a 100 *μ*m wide region of the epithelium.

**Figure 5.**
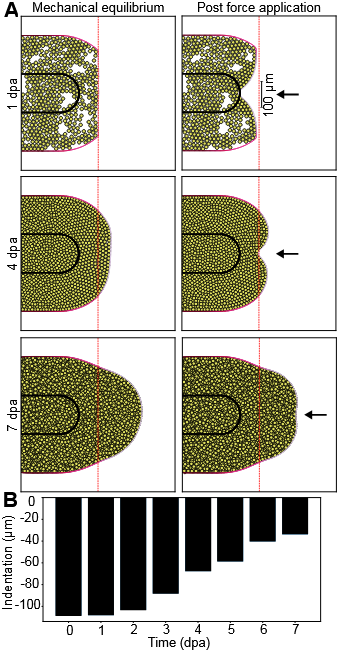
Quantification of composite tissue deformation to applied force reveals whole tissue stiffening at the transition from wound healing to regenerative blastema formation. (A) Simulated tissue deformations (across) due to applied force at 1, 4, and 7 dpa. (B) Measurements of limb blastema deformation for constant applied force across daily time point since amputation.

We quantify the in-plane deformation of the limb as a result of this externally applied force (Fig. 5B). We find that the amount of tissue deformation correlates negatively with the time since amputation, suggesting an overall stiffening of the blastema. This result reveals an emergent increase in blastema stiffness over time, despite local epithelial softening. In our model, this stiffening is the result of proliferation induced increases in mesenchymal cellular density. This suggests a composite tissue response that agrees with recent experimental findings in other model organisms [48].

### Model parameter calibration to blastema images reveals Wnt inhibition disrupts directed motility in the mesenchyme

Next, we turn our attention to the effect of complete Wnt inhibition. We and others have reported that the C59 inhibitor halts the formation of a blastema; yet, we did not find significant differences in either the overall number of proliferating cells or their spatial distribution within the blastema, as indicated by EdU staining (Fig. 1G). By evaluating the morphometrics of the simulated outgrowth (length, area, and aspect ratio), several hypotheses consistent with the C59 perturbation emerge: phase separation, preferentially oriented division along the PD or AP axes, and a proliferation gradient (Fig. 3A-C, Fig. S4). The absence of migration towards the distal tip is a shared characteristic across all these hypotheses.

To confirm this finding, we adapted a statistical inference method to determine the model parameter values (migration, proliferation rates) directly from the experimentally-obtained blastema shape (Fig. 6). The algorithm infers parameter values by minimizing the shape error, defined as the root mean squared error (RMSE) between the experimental and simulation-obtained blastema boundaries at 7 dpa, parameterized by the angular coordinate along the boundary: 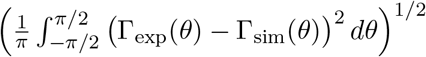 . Three parameters were determined using this method: (1) the total cell cycle duration 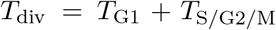 with 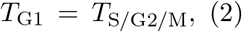 the proportion of mesenchyme cells, m, migrating towards the distal tip, and (3) the migration speed (*μ*). To handle stochastic noise in our shape error, we use a modified particle swarm optimization (PSO) algorithm. Particles representing candidate parameter sets (*T*_div_, *m, μ*) move through parameter space with velocities influenced by both their personal best position and their neighborhood’s best position. We employ the *lbest k* = 2 topology, where each particle’s neighborhood consists of its two nearest neighbors. This local topology performs better than global best topologies in multi-modal landscapes to avoid premature convergence and promote space exploration [49]. Particles are initialized within regions informed by preliminary simulations, and parameter values are bounded. To further promote exploration and avoid getting stuck in a local minimum of the shape error, particles are periodically given random velocity perturbations. These perturbations are manually applied when over 75% of the parameter space is unexplored prior to convergence.

**Figure 6.**
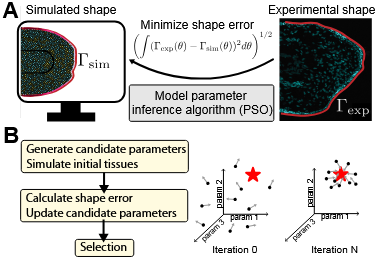
(A) Schematic of the pipeline for parameter estimation showing identification of model parameters that minimize RMSE between the simulated shapes and experimental shapes of blastema outgrowth. (B) Flowchart of the inference parameter estimation pipeline utilizing the particle swarm searching algorithm for the global minimum of the shape error.

We ran the inference pipeline for two different experimental curves: one obtained as an ensemble average of the epithelial boundary from control experiments and the other obtained from the ensemble average of the C59 treated samples. Motivated by our AFM measurements, the epithelial stiffness profile is fixed as locally softer, implemented in a region 400 *μ*m centered about the midline. Fig. 7A shows the resulting shape error at various parameter sets for control limb blastemas. A negatively sloped manifold with low RMSE is identified, which can be more clearly seen in the (*k*_div_, *m*) parameter space for a fixed migration speed of *μ* = 100 *μ*m/day (Fig. 7B). A similar manifold is observed for C59 (Fig. 7D-E), but with substantially lower values of *k*_div_ and *m*. This suggests that each suitable (*k*_div_, *m, μ*) pair must maintain a particular ratio to generate a blastema of a prescribed shape. This matched our intuition that the rates of motility and cell division could not be distinguished simply from the outgrowth shape, and further information would be needed to constrain this parameter inference problem.

**Figure 7.**
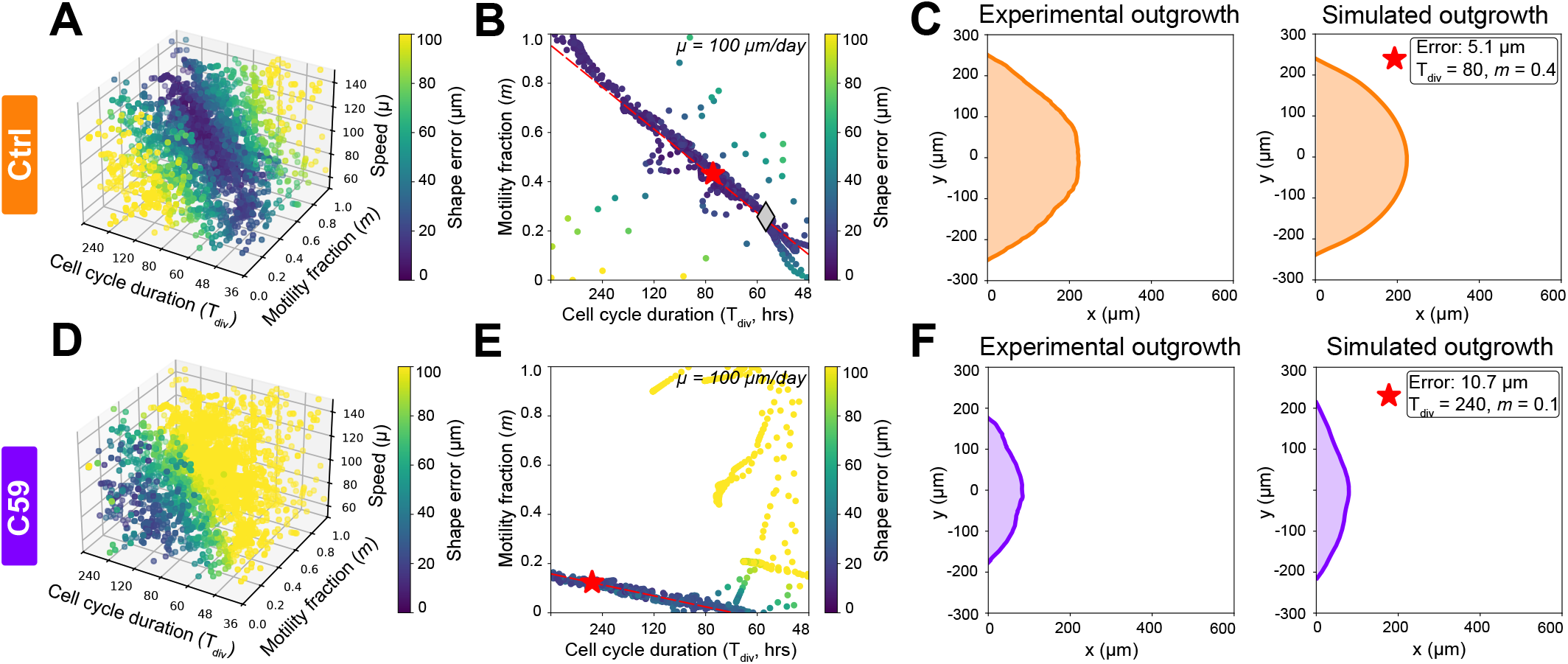
Fitting results of kinetic rates in hybrid agent-based model based on averaged blastema shape in control and Wnt inhibited limbs at 7 dpa. (A,D) Simulation parameter combinations (*k*_div_, *m, μ*) pairs colored by shape error for control (top) and C59 (bottom) limbs. (B,E) Select optimized parameter combinations along the plane *μ* = 100 *μ*m/day. Star indicates the parameter combination with lowest error, diamond refers to (*k*_div_ = 0.42, *m* = 0.3) as in Fig. 3. (C,F) Experimentally obtained (left) and best fit simulation (right) for control and C59.

Minimum shape error for control limbs was achieved with a cell cycle of roughly 80 hours and a 40% motility fraction of mesenchymal cells at a fixed speed of 100 *μ*m/day (Fig. 7C). In contrast, Wnt-inhibited limbs required a much longer cell cycle (over 240 hours) and a lower motility fraction (less than 15%) to minimize shape differences (Fig. 7F). This suggests that Wnt inhibition either reduces proliferation events or impairs the directional migration of mesenchymal cells toward the wound epithelium. Since our data show no changes in the number of S-phase cells under full Wnt inhibition (Fig. 1G), we conclude that Wnt signaling indeed controls the directional migration of cells into the blastema tissue. Lastly, when we repeated the results using an epithelial layer of uniform mechanical stiffness (but softer), the model produced a similarly shaped blastema as in control experiments when the cell cycle was over 80 hours, well above what has been estimated by others [18] (Fig. S6A). In contrast, for C59 to halt outgrowth, an even longer cell cycle (over 120 hours) was needed, along with a reduced fraction of distal-tip-directed motile cells (Fig. S6B).

## DISCUSSION

We present a hybrid agent-based modeling framework for forward simulating the formation of the accumulation stage blastema. A parameter inference pipeline is used to extract kinetic model rates of cellular dynamics underlying a particular shape. Our framework makes two important biological discoveries (Fig. 8) — (1) local softening of the epithelial layer post-wound healing at 7 dpa, and (2) directional migration of mesenchyme cells towards the wound epithelium/AEC requires Wnt signaling. AFM measurements of the regenerating epithelium confirm that the stiffness patterning persists, despite recent seemingly contradictory results at the whole tissue level [48, 50]. We explain this discrepancy via a composite multilayered tissue response: the model simultaneously recapitulates the stiffening of the whole limb with the softening of the outermost layer at the injury site. Cell proliferation results in an exponential increase in the number of cells and a consequential increase in volume fraction, all of which offer mechanical resistance to external tissue indentations.

**Figure 8.**
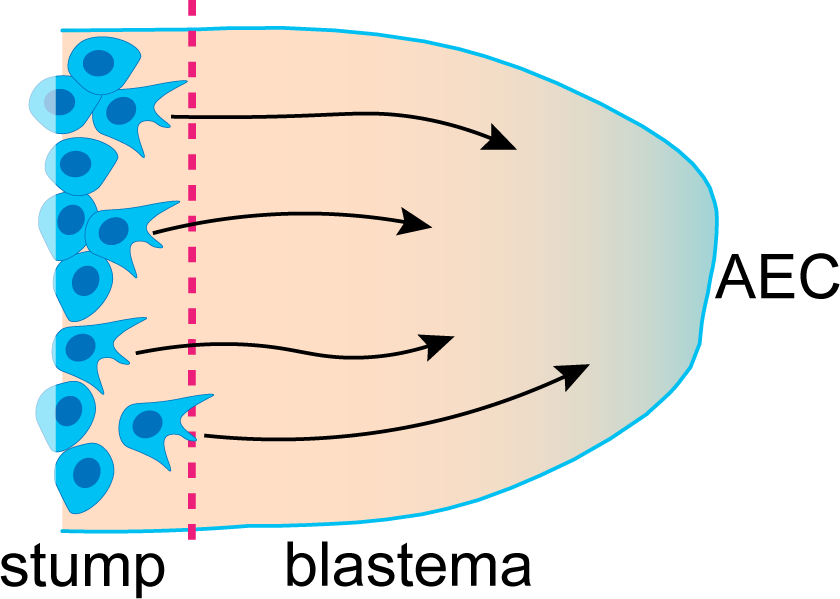
Graphical abstract. Model of limb blastema formation through local softening of the wounded epithelium/AEC and AEC-directed migration of mesenchyme cells controlled by Wnt signaling.

Mechanical patterning of a developing tissue’s surroundings directly and indirectly contributes to its final form, either by restricting or permitting deformation in a number of developing systems [51]. Expansion of the brain ventricle lumen during hindbrain morphogenesis is only possible through the softening of the surrounding epithelium, and in stiff mutants, this is impaired because the ventricle fails to enlarge [52]. In a multi-step process, mechanical patterning of surrounding tissues produced the vilification of the gut [33, 53]. Elongation of the *Drosophila* egg chamber is driven by the collective migration of follicular epithelial cells, which are guided by a surrounding extracellular matrix that is heterogeneously stiff [54, 55]. The cause of the observed local softening of the epithelium could be the result of a number of events, including EMT transitions and local matrix metalloproteinases (MMP) degradation of the basement membrane and the extracellular matrix in the mesenchyme [56], or induced fibroblast migration during wound healing [57].

Our method is not without limitations. First, our model is limited to a 2D cross-section and ignores out-of-plane dynamics and deformations. Second, cellular shape is ignored; however, future work will include extensions that can incorporate and/or validate experimental data on morphological differences. Third, although proliferation in the epithelial layer is not explicitly modeled here, there is continual migration of these cells into the wound epithelium/AEC. A more detailed model of the epithelium could account for these dynamics as well as provide a mechanistic understanding of the cellular dynamics (polarization, weakened cell-cell adhesion) that produce this apparent mechanical patterning of the epithelium. Lastly, we report that, without additional constraints, the parameter inference pipeline cannot circumvent parameter identifiability issues — for example, observing the net growth rate cannot distinguish between proliferation and apoptosis rates.

## Author contributions

C.C. and J.M. conceptualized and developed the research project. S.F. performed most of the experiments and data analysis with help from D.T.F. and X.W. A.B.-S. and C.C. developed the computational model with help from X.W. A.B.-S. ran and analyzed all computational simulations. C.C., S.F, and A.B.S. prepared the manuscript and figures with input from J.M. All authors edited the manuscript. C.C. and J.M. supervised the project.

## Acknowledgments

This work was supported in part by NSF DMS2209494 (C.C.) and NICHD R01HD099174 (J.M.). We thank the Institute of Chemical Imaging of Living Systems at Northeastern University for consultation and imaging support.

## METHODS

### Animal ethics statement

Animal experimental procedures were approved by IACUC. All d/d genotype axolotls (RRID:AGSC 101E; RRID:AGSC101L) were bred at Northeastern University or obtained from the Ambystoma Genetic Stock Center (RRID: SCR 006372).

### Animal experimental procedures

Juvenile white (d/d) axolotls, between 2.8 and 3.5 cm long in total length measured from snout to tail tip, were used for drug inhibition experiments. Animals were anesthetized using 1X benzocaine; a toe pinch was performed to ensure that the animals were properly anesthetized before amputations were conducted. Animals were bilaterally amputated through the distal stylopod directly above the elbow joint. During normal regeneration, animals were housed in housing water with DMSO, while the animals treated to inhibit regeneration were housed in housing water with 5 *μ*M Wnt-C59 (Tocris, Cat. No. 5148). Animals were left to regenerate for 7 days after amputation; drugged housing water was replaced every 2 days during the normal feeding and cleaning routine. An EdU pulse was performed 3 hours prior to limb collection; the animals were given an IP injection of EdU (10.57 *μ*g/kg body mass) to label cells that are currently in the S phase of the cell cycle.

### Histology

The collected forelimbs were immediately transferred to 4% paraformaldehyde (PFA) and fixed overnight at 4C. Samples were then cryoprotected in 30% sucrose in PBS at room temperature or at 4C overnight until they sank. Samples were embedded in optimal cutting temperature (OCT) compound and frozen at 80C. Cryosectioning was performed at a thickness of 16 *μ*m. Tissue sections were treated with DAPI to label the nuclei and Cytoliner, a lipophilic fluorescent dye (Biotium, #30134, 1:500), to label the cell membrane. Click-it chemistry was used to label cells in the S phase of the cell cycle. Following 1x SSCT washes, samples were incubated with Click-iT cocktail (888 *μ*L 1x TRIS buffered saline, 10 L 100 mM CuSO4, 2 *μ*L 500 mM azide- or alkyne-modified probes, and 100 *μ*L 0.3M sodium ascorbate) for 30 minutes in the dark. Samples were mounted with SlowFade Gold Antifade Mountant.

### Imaging

Images were taken on a Zeiss 880 confocal microscope using a 20x air objective. All acquisition parameters were kept consistent across various samples. Images were post processed using Zen Black prior to data analysis to stitch and perform maximum intensity projections across 10-14 z-plane sections with a step size of 1 *μ*m.

### Cell segmentation, cell cycle and morphological assessment

Images were analyzed using the ARIVIS software. Cells were segmented using CellPose-SAM on the DAPI channel; EdU fluorescence was segmented using thresholding. Parent-child analysis was used to group objects within 500 *μ*m of the distal tip of the limb; objects were grouped in 100 *μ*m sections across the entire limb, starting at the distal tip. Objects were also grouped by tissue type; we categorized them into the epithelium and mesenchyme. Each of these grouped objects was further pooled based on EdU fluorescence. Object measurements, both tissue wide and individual cells, were analyzed using MATLAB and GraphPad Prism. To determine the spatial distribution of cells in S phase, EdU positive cell counts were divided by the total cell count for each grouping of cells. The aspect ratio of each cell is calculated as the maximum height divided by twice the maximum width, such that a semicircle would have a unit aspect ratio.

### AFM measurements

#### Sample preparation

Axolotls measuring an average of 3 cm in total length underwent bilateral amputations of the forelimbs at the upper arm and were left to regenerate for 7 days. At 7 dpa, the limbs were collected and embedded in 8 % agarose in deionized water. Limbs were sectioned at 100 microns on a Leica VT1000 S Vibratome. Tissue samples were adhered to glass bottom dishes by applying krazy glue to the outer corners of the agarose sections; the dish was then filled with 0.8x PBS.

#### AFM measurements

Force curves were obtained by performing indentations using a cantilever with a triangular geometry and a 20 *μ*m Borosilicate glass particle attached (PT.BORO.SN.20 from NovaScan Technologies) with an estimated stiffness of 0.1 N/m. Cantilevers were manually calibrated before each experiment. Contact mode spectroscopy with a setpoint of 2V to probe the bottom of a sterile glass dish containing 0.8X PBS was used during manual calibration.

Measurements of injured and uninjured epithelium were performed using a setpoint of 0.5 nN and 2.5 nN. All measurements were conducted with an approach speed of 2 *μ*m/s, a retraction speed of 2 *μ*m/s, a contact time of 0 s, and a z length of 7.5 *μ*m.

#### Force curve analysis

Force curve analysis was conducted using Bruker provided JPK SPM data processing software version 8.1.61. Built in processing tools were used to determine the young’s modulus from the collected force curves. Data smoothing, baseline subtraction, contact point determination, vertical tip position, and elasticity fit were all used in series to determine the apparent Young’s modulus.

