## Supporting Information for "Model recapitulates regenerative limb blastema formation through local softening of the wounded epithelium"

#### SUPPLEMENTAL FIGURE LEGENDS:

**SUPPLEMENTAL FIGURE 1**: **Spatial quantification of cell morphology and cell cycle state in the regenerative blastema at 7 dpa.** (A) Schematic of 100 $\mu$m physical sections from the distal tip. Red line, amputation plane. Green line, 500 $\mu$m region. (B-C) Quantification of the area and aspect ratio of nuclei in the blastema tissue by 100 $\mu$m sections in each condition. (D) Quantification of EdU+ (S-phase) cells by 100 $\mu$m sections in each condition. $N = 5$ animals/condition except in (D) where $N = 6$ animals were used for C59. Dashed horizontal lines, average for the entire 500 $\mu$m region for control (orange) and C59 (purple). Mann-Whitney test, $* \, p < 0.05$.

**SUPPLEMENTAL FIGURE 2**: **Cell cycle implementation details.** (A) Runtime per time step scales linearly with the number of cells. Dashed red line, line of best fit. (B)-(C) Validation that the stochastic processes of proliferation match the expected ensemble continuum behavior both overall and when distinguished by phases. Dashed lines, ODE solution integrated numerically via Runge-Kutta methods.

**SUPPLEMENTAL FIGURE 3**: **Volume fraction estimates in experiments and simulations.** Evolution of mesenchyme volume fraction over time in a simulation with local softening of the epithelium. Inset, sample of blastema tissue collected at 7 dpa and stained with Cytoliner membrane marker in purple. Dashed red line, amputation plane.

**SUPPLEMENTAL FIGURE 4**: **Shape error for simulated outgrowth hypotheses.** The root mean squared error (RMSE) between the experimental and simulation-obtained blastema boundaries at 7 dpa for (A) normal and (B) C59/Wnt-inhibited limbs for all simulated hypotheses.

**SUPPLEMENTAL FIGURE 5**: **Effect of various spatial patterning of epithelium on blastema outgrowth morphometrics.** Contour plots of simulated blastemas at 7 dpa illustrating how distinct assumptions about epithelial spatial stiffness gradients influence tissue morphology. (a) The entire distal limb region is mechanically softer. (b) A localized region of reduced mechanical stiffness (400 $\mu$m wide) is centered at the distal tip, with additional randomly distributed softer patches. (c) The same localized 400 $\mu$m is shifted by 200 $\mu$m to the right of the distal tip center, generating an asymmetric spatial gradient. Epithelium contours are shown, with lighter shading indicating lower mechanical stiffness. Representative trajectories of 30 cells over 7 dpa are overlaid to illustrate mechanistically driven migration patterns.

**SUPPLEMENTAL FIGURE 6**: **Fitting results of kinetic rates in hybrid agent-based model for spatially uniform epithelial layer.** Resulting shape error for simulation parameter combinations $(k_{\text{div}}, m)$ along the plane $\mu = 100$ $\mu$m/day for (A) control and (B) C59 blastemas. Star indicates the parameter combination with lowest error.

#### SUPPORTING MOVIES:

**Movie 1**: 2D simulation of blastema outgrowth from a flat wounded epithelium with randomly oriented division planes during proliferation events, apoptosis, and Brownian motion. G1 phase (rest), yellow cells. S/G2/M phase (dividing), cyan cells.

**Movie 2**: 2D simulations of blastema outgrowth under the assumption of (top) distal tip directed migration of 30% of mesenchymal cells and (bottom) phase separation, together with 50% of cells undergoing distal tip directed migration. A Side-by-side comparison of these two hypotheses, with and without local softening of the epithelium.

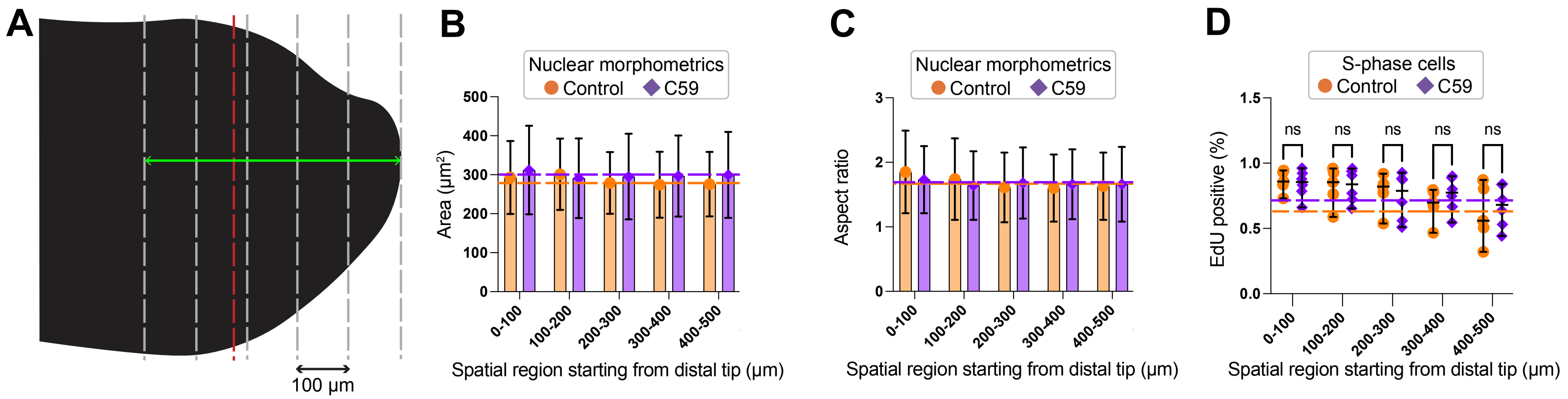
A
100 μm
B
Nuclear morphometrics
Control
C59
Area (μm²)
500
400
300
200
100
0
0-100
100-200
200-300
300-400
400-500
Spatial region starting from distal tip (μm)
C
Nuclear morphometrics
Control
C59
Aspect ratio
3
2
1
0
0-100
100-200
200-300
300-400
400-500
Spatial region starting from distal tip (μm)
D
S-phase cells
Control
C59
EdU positive (%)
1.5
1.0
0.5
0.0
ns
ns
ns
ns
ns
0-100
100-200
200-300
300-400
400-500
Spatial region starting from distal tip (μm)

**Figure S1**

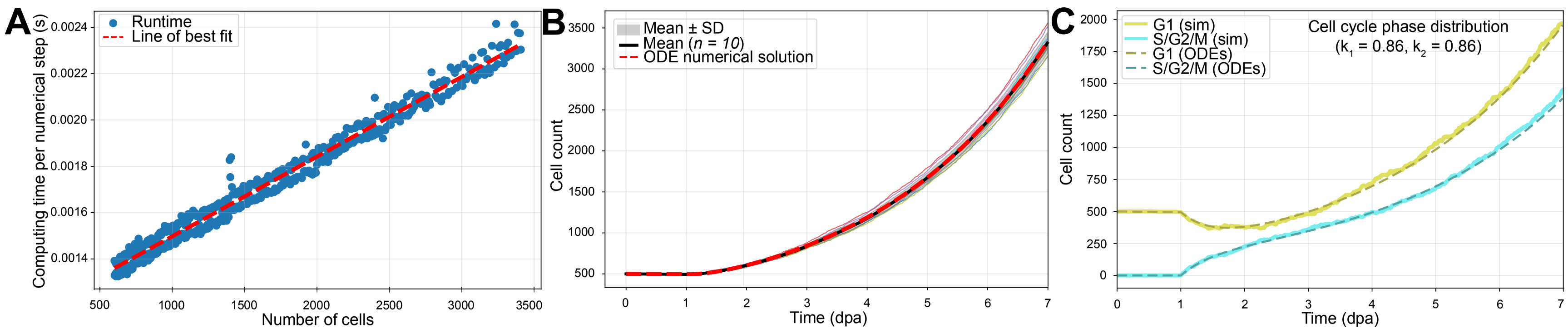
A
Runtime
Line of best fit
Computing time per numerical step (s)
0.0024
0.0022
0.0020
0.0018
0.0016
0.0014
500
1000
1500
2000
2500
3000
3500
Number of cells
B
Mean ± SD
Mean (n = 10)
ODE numerical solution
Cell count
3500
3000
2500
2000
1500
1000
500
0
1
2
3
4
5
6
7
Time (dpa)
C
G1 (sim)
S/G2/M (sim)
G1 (ODEs)
S/G2/M (ODEs)
Cell cycle phase distribution
(k1 = 0.86, k2 = 0.86)
Cell count
2000
1750
1500
1250
1000
750
500
250
0
0
1
2
3
4
5
6
7
Time (dpa)

**Figure S2**

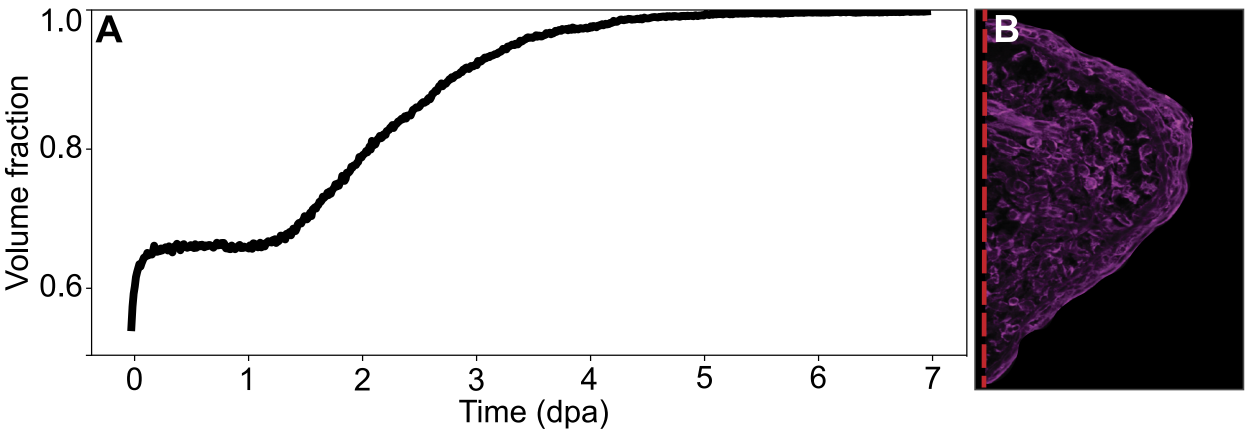


**Figure S3**

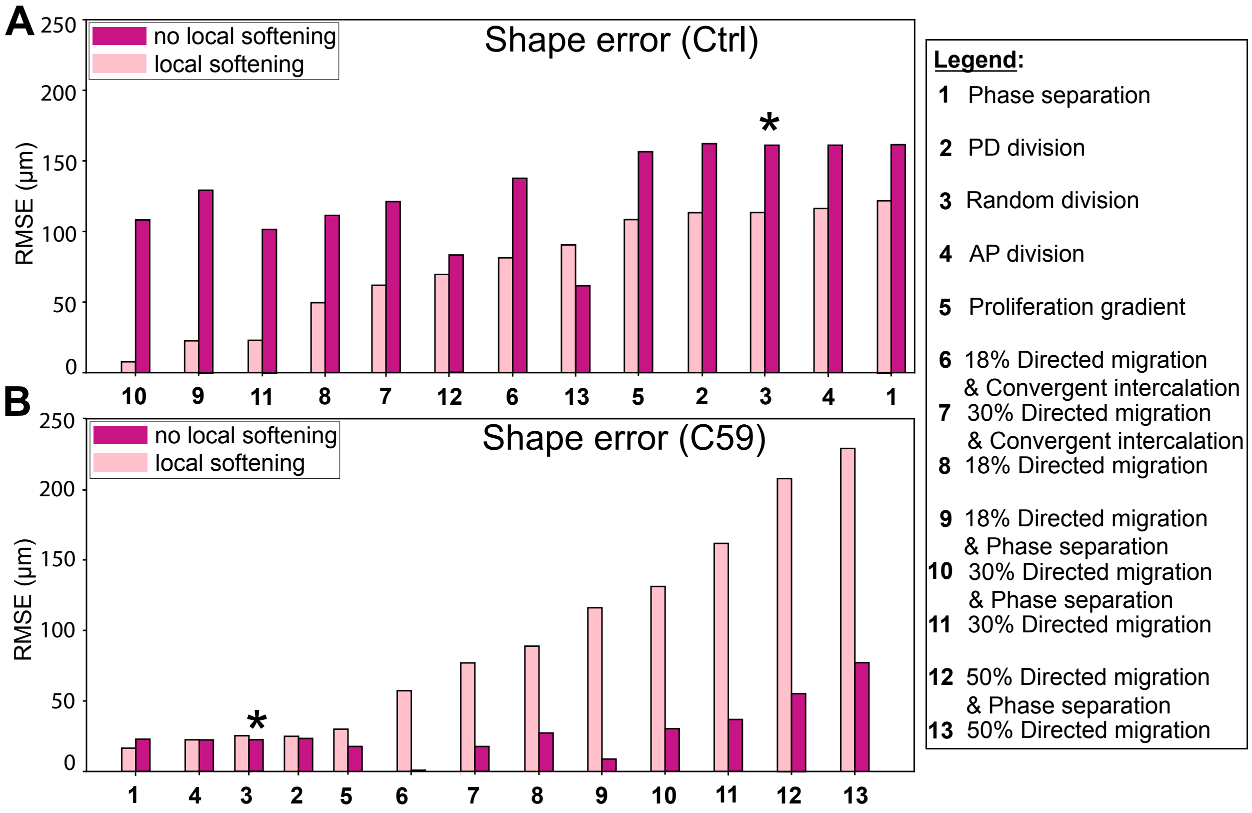
A
Shape error (Ctrl)
no local softening
local softening
RMSE (μm)
0
50
100
150
200
250
10 9 11 8 7 12 6 13 5 2 3 4 1
*
B
Shape error (C59)
no local softening
local softening
RMSE (μm)
1 4 3 2 5 6 7 8 9 10 11 12 13
Legend:
1 Phase separation
2 PD division
3 Random division
4 AP division
5 Proliferation gradient
6 18% Directed migration & Convergent intercalation
7 30% Directed migration & Convergent intercalation
8 18% Directed migration
9 18% Directed migration & Phase separation
10 30% Directed migration & Phase separation
11 30% Directed migration
12 50% Directed migration & Phase separation
13 50% Directed migration

**Figure S4**

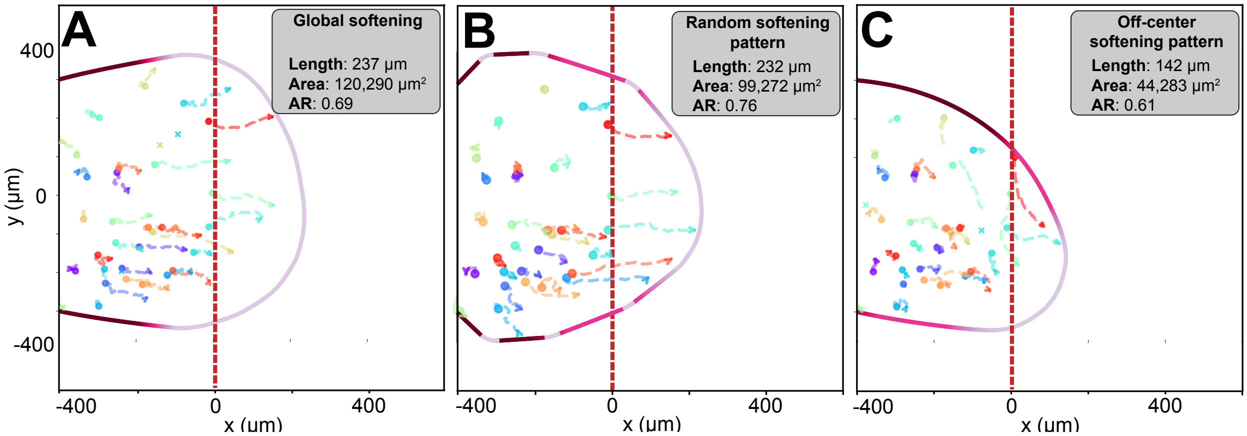
A
Global softening
Length: 237 µm
Area: 120,290 µm²
AR: 0.69
B
Random softening pattern
Length: 232 µm
Area: 99,272 µm²
AR: 0.76
C
Off-center softening pattern
Length: 142 µm
Area: 44,283 µm²
AR: 0.61
400
0
-400
y (µm)
-400
0
400
x (µm)

**Figure S5**

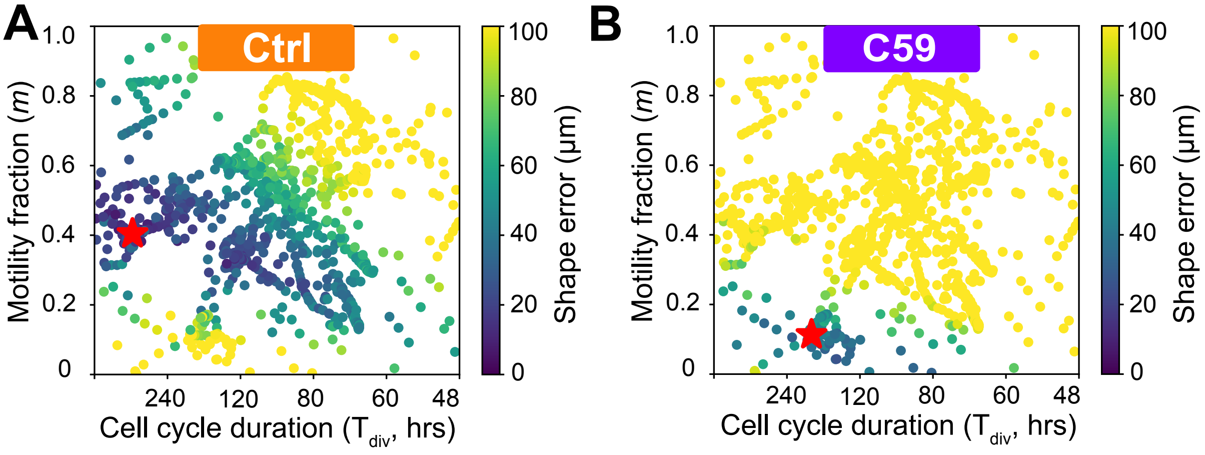
A
Ctrl
Motility fraction (m)
1.0
0.8
0.6
0.4
0.2
0
240
120
80
60
48
Cell cycle duration (T_div, hrs)
Shape error (μm)
100
80
60
40
20
0
B
C59

**Figure S6**

### Supporting Information for "Model recapitulates regenerative limb bud formation through local softening of the injured epithelium"

Samantha Finkbeiner [a] Ansa Brew-Smith [b] Xueqing Wang [b]
Dzi Tsiu Fu [a] James Monaghan [a,b] Calina Copos [a,b]

#### A. Numerical implementation of hybrid agent-based model

The code for this computational framework was developed in Python, and it is available on GitHub: `https://github.com/CoposGroup/Regen_ABM`. The equations of motion for the mesenchyme and epithelium in Eqs. (1)-(2) assume that cellular dynamics occur in the overdamped regime, which is a valid modeling assumption in most cellular biological systems. Each Langevin equation includes a constant damping term. Equations are numerically solved using an explicit Forward Euler integration scheme. The positions of mesenchymal cells and epithelium nodes are stored as NumPy arrays of size $(20N_{\mathrm{M}}, 2)$ and $(N_{\mathrm{E}}, 2)$, respectively, with all calculations performed in-parallel via vectorized operations using the Numba just-in-time compiler. A random dispersion of cells was used as the initial condition, with $N_M = 500$ mesenchymal cells placed in a region that represents the limb mesenchyme behind a flat amputation plane (Fig. 2B). The amputated limb domain is 600 $\mu$m wide by 400 $\mu$m long. Parameters for the initial shape of the limb tissue are taken from images of different 2D cross-sections by our group (Fig. 1C). Simulation was run for $1.4 \times 10^6$ time steps, with a time step size of $\Delta t = 5 \times 10^{-6}$ day, which corresponds to 7 days. To keep track of apoptosis events, a cell's existence is encoded into position arrays using `NaN` values. Forces between discrete agents (cell-cell, cell-epithelium) are computed using a cell-list algorithm, reducing the computational complexity from quadratic to linear scaling with the number of cells (Fig. S2A). Elastic contractile forces within the epithelium are computed using forward and backward finite difference matrices to quickly update edges between nodes. To prevent cells from escaping the blastema region, a ray-casting algorithm is used to detect escaped cells and place them inside the blastema at the closest epithelial node via a coarse-to-fine algorithm search. Stochastic processes such as cell proliferation, apoptosis, and drift-diffusion are computed in parallel through vectorized NumPy operations.

#### B. Justification of model parameters

Parameter values are provided in Table S1.

**Cell cycle length and apoptosis rate.** In our model, we find that blastema size scales strongly with cell count; namely, increased proliferation leads to more growth. Here, the total cell cycle is approximately 56 hours (matching literature values between 40-53 hours). Stochastic cell cycle dynamics were validated against an ensemble-averaged behavior in Fig. S2B. In the absence of additional information at these early regenerative time points, we divided the cell cycle into equal periods for two phases: G1 (rest) and S/G2/M (dividing). However, we do explicitly model the two distinct phases with distinct transition probabilities that match the ensemble behavior provided in

| Description | Parameter | Value (unit) |
|---|---|---|
| Mesenchyme repulsion strength | $k_{\mathrm{M,rep}}$ | 30.0 (a.u.) |
| Mesenchyme adhesion strength | $k_{\mathrm{M,adh}}$ | 0.15 (a.u.) |
| Mesenchyme–epithelium repulsion constant | $k_{\mathrm{ME,rep}}$ | 80.0 (a.u.) |
| Mesenchyme–epithelium adhesion constant | $k_{\mathrm{ME,adh}}$ | 0.01 (a.u.) |
| Random motion variance coefficient | $\sigma_{\mathrm{rm}}$ | $1\ \mu\mathrm{m}/\sqrt{\mathrm{day}}$ |
| Directed motion variance coefficient | $\sigma_{\mathrm{dm}}$ | $1\ \mu\mathrm{m}/\sqrt{\mathrm{day}}$ |
| Directed motion mean coefficient | $\mu_{\mathrm{dm}}$ | $100\ \mu\mathrm{m}/\mathrm{day}$ |
| Cell radius | $d_0$ | $20\ \mu\mathrm{m}$ |
| Frictional drag coefficient | $\xi$ | 1.5 (a.u.) |
| Cell apoptosis rate | $k_{\mathrm{death}}$ | $0.01\ \mathrm{day}^{-1}$ |
| Mean G1–phase duration | $T_{\mathrm{G1}}$ | 28 hours |
| Mean S/G2/M–phase duration | $T_{\mathrm{S/G2/M}}$ | 28 hours |
| Epithelial stiffness | $\kappa$ | 75, 150 (a.u.) |
| Simulation time step | $\Delta t$ | $5 \times 10^{-6}$ day |

**Table S1: Table of model parameters** used in blastema formation in axolotl limb regeneration. Time-related quantities are expressed in real time (hours or days).

Eqs. (4)-(5) (Fig. S2C). In our model, we use an apoptosis rate $k_{\mathrm{death}} = 0.01$ per day, which is in agreement with Mescher et al., who used similar-sized animals to measure 1-2% apoptotic cells in control and 4% in denervated limbs.

**Cell migration speed:** While estimates of migration speed in the early regenerative blastema have yet to emerge, a simple calculation was used to approximate this model parameter. Assuming a quiescent period of 24-48 hours for wound healing and closure, we take the distance from the end of the mesenchyme to the amputation plane at 7 dpa; that is our maximum migration over time. If we assume cells move in the last move to the base of the blastema, then that corresponds to 500 $\mu$m over 5-6 days or roughly 80-100 $\mu$m/day. For our forward simulations, we take the upper limit of 100 $\mu$m/day for distal tip directed migration speed. Lastly, mesenchymal cells also undergo random Brownian motion. Using mean squared displacement curves, we validated that our random Brownian motion corresponds to an effective diffusion coefficient of 0.5 $\mu\mathrm{m}^2$/day. To obtain the random motion variance coefficient, we use the Einstein relation in 2D, $\mathrm{MSD} = \langle |\boldsymbol{r}(t) - \boldsymbol{r}(0)|^2 \rangle = 2nDt$, for $t = \Delta t$ and $n = 2$:

$$\begin{aligned}
\mathrm{MSD} &= 2(2)D\Delta t \\
\mathrm{Var}(x) + \mathrm{Var}(y) &= 4D\Delta t \qquad (\mathrm{Var}(x) = \mathrm{Var}(y) = \sigma_{\mathrm{rm}}^2 \Delta t) \\
2\sigma_{\mathrm{rm}}^2 \Delta t &= 4D\Delta t \\
\frac{\sigma_{\mathrm{rm}}^2}{2} &= D \quad (\text{plug in real values, } \sigma_{\mathrm{rm}} = 1\ \mu\mathrm{m}/\sqrt{\mathrm{day}}) \\
0.5\ \mu\mathrm{m}^2/\mathrm{day} &= D.
\end{aligned}$$

**The coupled effect of varying proliferation and apoptosis rates.** Only the difference between proliferation and death rates has meaningful effects on blastema formation in this model, as opposed to individual $T_{\mathrm{S/G2/M}}$, $T_{\mathrm{G1}}$, and $k_{\mathrm{death}}$ values. Unsurprisingly, the size of the blastema outgrowth scales with cell count; more proliferation and fewer apoptosis events lead to the largest outgrowth. Specifically, with local softening present, our model can reproduce a blastema with morphology similar to experiments without distal-biased cell migration, provided that the proliferation rate and repulsion constant $k_{\mathrm{M,rep}}$ are sufficiently large (see below). This scenario would correspond to a total cell cycle that is $T_{\mathrm{G1}} + T_{\mathrm{S/G2/M}}$ of 38 hours long (compared to the estimated literature values of 40-53 hours).

**The effect of varying mesenchyme repulsion strength.** The cell-cell repulsion constant within the mesenchyme $k_{\mathrm{M,rep}}$ has a significant impact on the resulting shape. When $k_{\mathrm{M,rep}}$ is large enough, blastema shape deformations are driven almost entirely by repulsion, with cell migration having negligible effects. Even with high repulsion, local epithelium softening is still needed to produce outgrowth comparable to the experiments. Low repulsion values result in high cell density in the mesenchyme and minimal outgrowth. Increasing cell diameter $d_0$ has an effect similar to increasing repulsion strength. To inform our numerical choice for the repulsion constant, we calculate the volume fraction of mesenchymal cells in a 7 dpa limb sample stained with a cell membrane marker (Cytoliner, a lipophilic fluorescent dye) (Fig. S3). We obtain an estimate of 85-90% occupancy for the mesenchymal outgrowth and 89-92% for the entire outgrowth tissue (depending on thresholding conditions). Using a value for the repulsion constant in Table 1, we find that the volume fraction is initially around 60% and evolves to 99% by 7 dpa.

**The effect of varying epithelium stiffness.** Fig. S5 compares quantitative blastema outgrowth metrics under different epithelial stiffness patterning assumptions. Uniform distal softening (Fig. S5A) reproduces the experimental outgrowth length and area but yields a reduced aspect ratio (0.69), producing a shorter, wider morphology rather than the characteristic cone shape, indicating that global softening is insufficient. Introducing a localized distal deformable region with additional soft patches (Fig. S5B) reduces the outgrowth area but results in a more cone-shaped morphology, as indicated by the aspect ratio. Lateral displacement of the localized soft region (Fig. S5C) further decreases the outgrowth size, demonstrating that both localization and spatial alignment of epithelial softening are required to restore the correct blastema morphology and growth.

To model the attachment of the limb to the animal body, an additional force constrains lateral deformations of the epithelium proportional to $\kappa^2$; that is, $\mathbf{F}_{\mathrm{lat}} = -k_{\mathrm{lat}}\kappa^2(y - y_0)$, where $y$ refers to the AP (vertical) coordinate of the epithelium node, and $y_0$ is the AP coordinate in the reference configuration. The parameter $k_{\mathrm{lat}}$ is small enough that $\mathbf{F}_{\mathrm{lat}}$ is negligible for $\kappa < 100$.

#### C. Modeling details of potential mechanisms for outgrowth

Implementation details for all hypotheses explored in Figs. 3 and S2 are provided below. The numbering in the parentheses matches the legends in both figures.

**Phase Separation (1).** Similar to mechanisms proposed in limb bud development, we assume that cells near the distal tip are more 'motile' compared to the rest of the blastema and limb tissue. The 'unjammed' (motile) cells are defined to be in a small region of length 180 $\mu$m near the distal tip, as shown in (Fig. 2B). All cells outside this region are 'jammed' (stuck). A cell's status can be reassigned as jammed or unjammed based on their position relative to this distal tip defined region. Jammed cells do not undergo Brownian (or drift) motion, while unjammed cells have $\sigma_{\mathrm{rm}}$ increased by a factor of 10. When directed cell migration is combined with phase separation, only unjammed cells can undergo directed migration.

**Oriented Cell Division Angle (2,3,4).** The placement of daughter cells could be random (3), placed at a uniformly chosen angle with respect to the mother cell, or oriented along either the PD (2) or AP (4) anatomical axes:

$$\theta_{\mathrm{PD}} \sim N\big(0, (\frac{\pi}{6})^2\big), \tag{S1}$$

$$\theta_{\mathrm{AP}} \sim N\big(\frac{\pi}{2}, (\frac{\pi}{6})^2\big), \tag{S2}$$

where $\theta_{\mathrm{PD}}$ and $\theta_{\mathrm{AP}}$ denote the angles in the cases of proximodistal oriented division and antero-posterior oriented division, respectively. Furthermore, for both cases, there is a 50% probability that the angle will be flipped by $\pi$ radians to ensure that daughter cells can appear on either side of their mother.

**Proliferation Gradient (5).** To simulate a proliferation gradient, we first initialize a regulatory front at the amputation plane. Starting at 48 hours post amputation, allowing for wound closure, a regulatory front moves 50 $\mu$m proximal to the amputation plane each day. Cells distal to the regulatory front have their $T_{\mathrm{G1}}$ reduced by 50% (18 hours), and cells proximal to the front have their $T_{\mathrm{G1}}$ increased by 50% (36 hours). This creates more cell proliferation near the distal tip. The regulatory front is meant to mimic the presence of morphogen signals, localized at the AEC, that spatially diffuse each day.

**Directed Cell Migration (6-13).** Migrating cells are subject to directed motion increments at each time step, distributed normally with a nonzero mean $\mu_{\mathrm{dm}}$ and a noise factor $\sigma_{\mathrm{dm}}$:

$$\eta_t^{\mathrm{dm}} \sim N(\mu_{\mathrm{dm}}\mathrm{d}t, \sigma_{\mathrm{dm}}^2\mathrm{d}t). \tag{S3}$$

$\eta_t^{\mathrm{dm}}$ is applied only in the direction of the distal tip, and Brownian motion $\eta_t^{\mathrm{rm}}$ is applied along both spatial directions. The percentage of migrating cells is defined during initialization, and any newly created daughter cells become migratory based on the migration percentage. Directed persistent migration is restricted to cells located distal to a regulatory front. Starting at 48 hours post amputation, allowing for wound closure, the regulatory front moves 50 $\mu$m proximal to the amputation plane each day. As mentioned above, the regulatory front is meant to mimic the presence of a diffusive morphogen localized at the AEC.

**Convergent Intercalation (6-7).** Inspired by mechanisms proposed in limb bud development, we consider the hypothesis of convergent intercalation. A percentage of cells is determined a priori to be identified as intercalating cells; intercalating cells are subject to directed motion increments oriented towards the proximodistal axis, stopping near the center where the bone would be located in the limb. Intercalating cells move according to the same $\mu_{\mathrm{dm}}$ and noise factor $\sigma_{\mathrm{dm}}$ as those of directed migration in Eq. (S3).
